# Semantic feature production norms for manipulable objects

**DOI:** 10.1101/2023.04.24.537452

**Authors:** Daniela Valério, Akbar Hussain, Jorge Almeida

**Author notes:** Correspondence should be sent to: Jorge Almeida, Proaction Laboratory, Faculty of Psychology and Educational Sciences, Rua do Colégio Novo, Coimbra 3000-115, Portugal.

## Abstract

Feature generation tasks and feature databases are important for understanding how knowledge is organized in semantic memory, as they reflect not only the kinds of information that individuals have about objects but also how objects are conceptually parse. Traditionally, semantic norms focus on a variety of object categories, and, as a result, have a small number of concepts per semantic category. Here, our main goal is to create a more finely-tuned feature database exclusively for one category of objects – manipulable objects. This database contributes to the understanding of within-category, content-specific processing. To achieve this, we asked 130 participants to freely generate features for 80 manipulable objects, and another group of 32 participants to generate action features for the same objects. We then compared our databases with other published semantic norms and found high structural similarity between them. In our databases, we calculated the similarity between visual, functional, encyclopedic, and action feature types. We discovered that objects were grouped in a distinctive and meaningful way according to feature type. Finally, we tested the validity of our databases by asking three groups of participants to perform a feature verification task while manipulating production frequency. Our results demonstrate that participants can recognize and associate the features of our databases with specific manipulable objects. Participants were faster to verify high-frequency features than low-frequency features. Overall, our data provide important insights into how we process manipulable objects and can be used to further inform cognitive and neural theories of object processing and identification.

## Background

We are constantly recognizing and discriminating a large number of objects, and we do it efficiently even when perceiving them from different viewpoints or when particular features are hidden from view. This raises a question: how do we recognize a simple object, such as a glass, despite large variations in shapes, colors, materials, and viewpoints? And how do we distinguish two similar objects, such as a glass and a cup? One important avenue for understanding how we do this relates to understanding the kinds of information we hold about these objects when perceiving, identifying, and distinguishing them – for instance, the shape and size of a glass, or the presence of a handle on a cup. In this study, we contribute to this debate by creating a comprehensive feature database specific to a single category (i.e., manipulable objects) and making it publicly available. This database will hopefully shed new light on how we efficiently recognize and distinguish objects.

Semantic norms (i.e., databases containing features produced by participants) have been used in the last few decades to create experimental stimuli, as well as to develop and test theories of semantic representation and object recognition in healthy (e.g., Taylor, Devereux, & Tyler, 2011; Warrington & Shallice, 1984) and brain-damaged individuals (Capitani, Laiacona, Mahon, & Caramazza, 2003; Duarte, Marquié, Marquié, Terrier, & Ousset, 2009; Marques, Raposo, & Almeida, 2013; Perri, Zannino, Caltagirone, & Carlesimo, 2012). These norms have improved our understanding of how information is represented and processed in the brain, both at a neural level (e.g., Kivisaari et al., 2019), and in a temporal perspective (e.g., Clarke, Taylor, Devereux, Randall, & Tyler, 2013). The most widely used set of semantic norms were created by McRae and collaborators (2005) and by Devereux and collaborators (2014), who collected features for a large number of living and non-living things, using 30 participants per concept. Similar norms were collected by numerous research groups in different countries and languages (de Deyne et al., 2008; Montefinese, Ambrosini, Fairfield, & Mammarella, 2013; J. Vivas, Vivas, Comesaña, Coni, & Vorano, 2017), showing high similarities in features lists across languages (Kremer & Baroni, 2011; Vivas et al., 2020). These norms have been used to study differences between living and non-living things (Goldberg, Perfetti, & Schneider, 2006; Randall, Moss, Rodd, Greer, & Tyler, 2004), and have led to proposals regarding putative differences in feature types and feature configurations between semantic categories (Cree & McRae, 2003; Garrard, Lambon Ralph, Hodges, & Patterson, 2001). However, these norms only include a few concepts per semantic category, making it difficult to gain a more specific understanding of how content-specific representations stripped of inter-category variability are created and stored cognitively and (perhaps) neurally.

In this study, we create a more detailed database to enhance our understanding of content-specific processing within a category. We chose the category of manipulable objects, as a case study, because they possess a set of interesting properties: 1) these objects have particular functions that they fulfill; 2) they require typical motor programs associated with their use; 3) they have specific structural features (e.g., being round); 4) they are used in specific contexts, and 5) we perceive and interact with them constantly. Moreover, patients can experience severe impairments in recognizing manipulable objects, compared to other categories of stimuli (e.g., Hillis & Caramazza, 1991; Moss & Tyler, 2000).

Here, we present two databases containing an extensive list of features for 80 manipulable objects. To assess the reliability of our databases, we compared them with the feature norms collected by McRae (McRae, Cree, Seidenberg, & McNorgan, 2005) and CSLB (Devereux, Tyler, Geertzen, & Randall, 2014). Our databases are highly comparable to these established multicategory norms. Finally, we used a feature verification task to demonstrate the validity of our features and databases, which showed that feature production frequency affects feature verification accuracy and reaction times.

## Experiment 1 – Feature generation and comparison with McRae and CSLB norms

Our data collection encompassed two stages. In the first stage, we asked participants to freely generate features for the objects. In the second stage, another group of participants freely generated features related to how one physically uses these objects (i.e., features related to object-specific action). We specifically asked participants for these action features because they are an important aspect of our understanding of manipulable objects, as demonstrated by many studies with apraxic patients (Almeida et al., 2018; Buxbaum & Saffran, 2002; Garcea, Dombovy, & Mahon, 2013; Valério et al., 2021), and because they can be more difficult to describe spontaneously. All objects were presented visually in a word format (see Table 1 of Supplementary Material for an overview of the 80 objects).

**Table 1:**
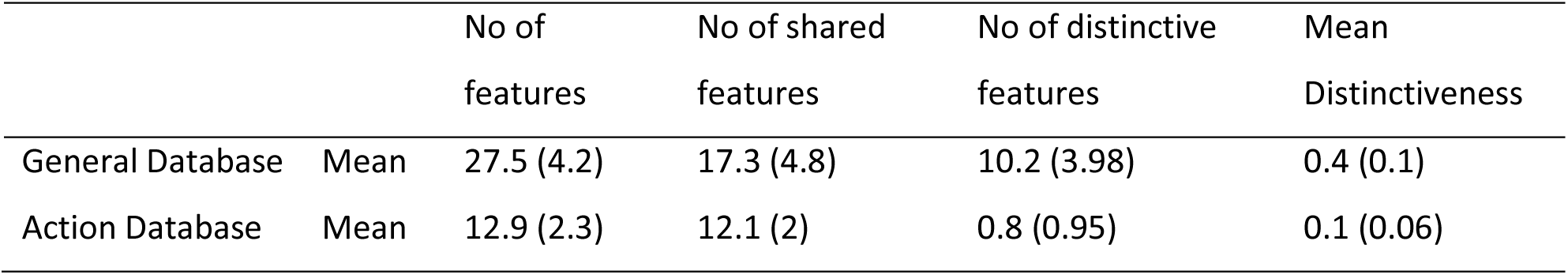
Mean and standard deviation (SD) of the Number of Features per object, Number of shared features, Number of distinctive features, and Mean of Distinctiveness for General and Action databases.

To compute our norms, we calculated measures of informativeness, including production frequency, distinctiveness, similarity between objects, category, and familiarity (see the Results section of Experiment 1). Using these measures, we then compared our databases to those of McRae and CSLB norms.

## Methods

### Participants

One-hundred and thirty participants (112 women), with an average age of 18.76 years (ranging from 18 to 35) listed features for 80 manipulable objects. Participants were instructed to list all the features that came to their minds. Another 32 participants (no overlap with those from the general feature production experiment) were asked to describe how to grasp and manipulate the same 80 objects using simple sentences. These participants included 29 women. All participants were right-handed except two, with an average age of 20.19 years (ranging from 18 to 37). To test how familiar participants were with the 80 objects, we asked an additional 50 subjects (43 women) to rate their familiarity with each object. The average age of this group of participants was 20.12 (ranging from 18 to 31).

All participants were native Portuguese speakers. They were first-year students of the University of Coimbra and received a course credit for their participation. All participants provided written informed consent for the study according to the Ethics committee of the Faculty of Psychology and Educational Sciences, University of Coimbra.

### Data Collection

Participants met with an experimenter who explained the experiment, and written consent was obtained from them to participate in the study. For the General Database, participants received a file (by email) that contained instructions for filling out the document and a page for each of the 80 objects presented in randomized order (per participant). Participants listed features in Portuguese on each page. We instructed our participants not to search the objects on the internet and to move on to the next object after listing features for that object without returning to previous ones. After completing the task, participants sent the completed experiment back by email.

For the Action database and Familiarity tasks, we used an online survey. Participants received a link with the instructions, and the objects were presented individually (per participant). After completing the task, participants submitted it on the survey webpage. In the Action Database, we followed the same procedures as the main experiment, but this time we asked participants to describe how to grasp and manipulate each of the 80 objects. For the Familiarity task, participants rated their familiarity on a 9-point Likert scale, with nine indicating “extremely familiar”. See Figure 1A for an example of and object and features generated in the General and Action databases in response to that object.

**Figure 1:**
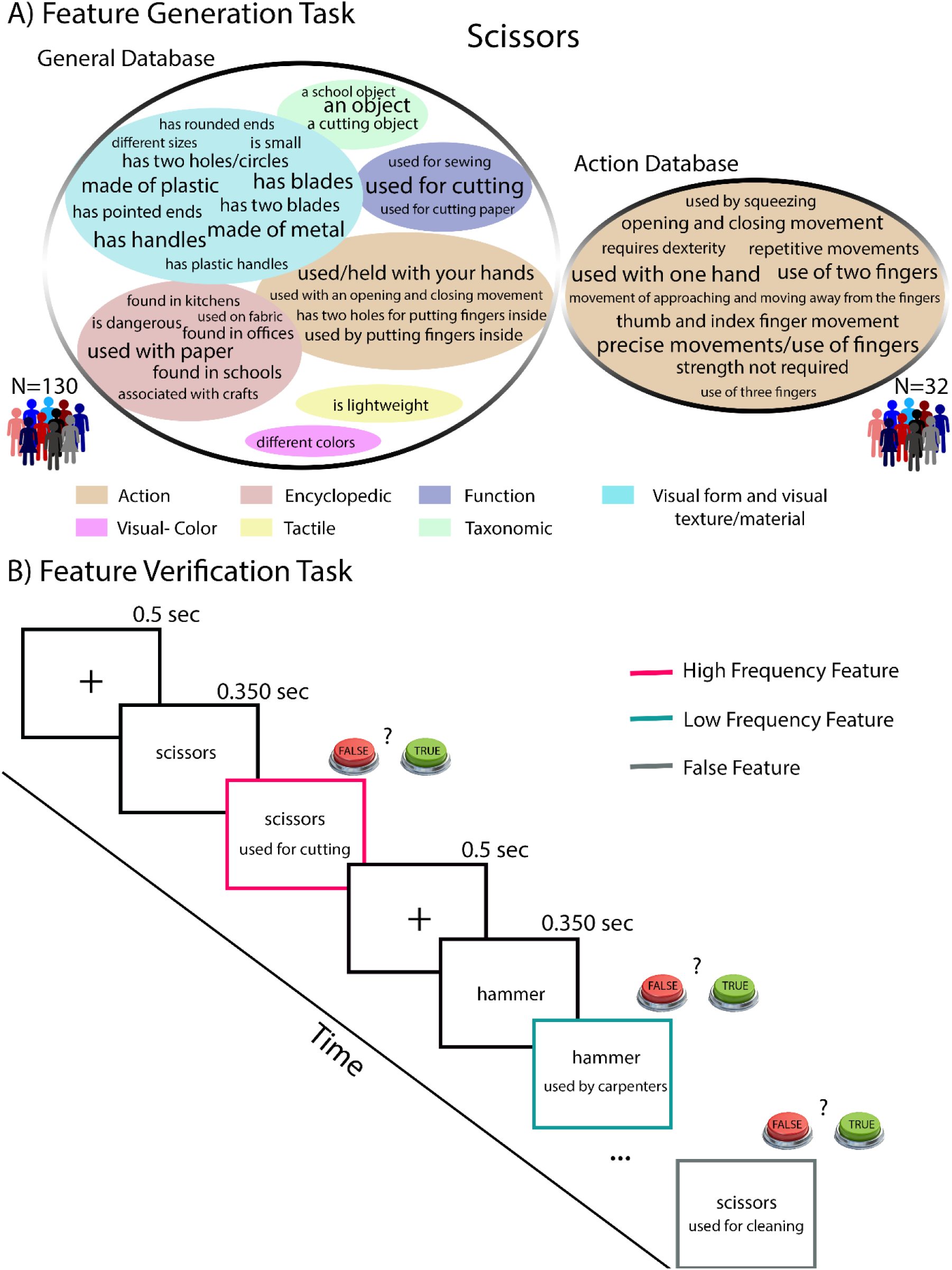
The two experiments A) Feature Generation Task, and B) Feature Verification Task. A) Example of features generated to the object scissors for both general and action databases, the font size represent production frequency, and features are clustered based on the category that they belong to; B) Experimental design for the feature verification task with three examples: High True Frequency Feature, Low True Frequency Feature and False Features.

### Comparison with McRae and CSLB norms

The McRae (McRae et al., 2005) and CSLB (Devereux et al., 2014) norms are perhaps the most widely used feature norms. It is therefore important to determine the degree of comparability between our databases and these norms. We found 23 manipulable objects in common between our databases and these norms. Those 23 objects are analyzed here in depth to compare our databases with those from the literature.

In our analysis, we calculated the number of features, number of shared features, and number of distinctive features for the General and the Action databases. We also collected these variables for the selected items from McRae and CSLB norms. To compare the similarity between the 23 objects, we used cosine similarity. We first organized rows and columns of the similarity matrices in alphabetical order to compare the pattern of shared features in both the General database and the McRae and CSLB norms (Fig. 2). Then, to assess the interpretability of the semantic similarity between the 23 objects, we used complete-linkage hierarchical clustering to organize objects into clusters (See Fig. S1 of the Supplementary Material) based on the data from the different databases. Additionally, we calculated the distribution of features across the different feature types (See Table 3).

**Figure 2:**
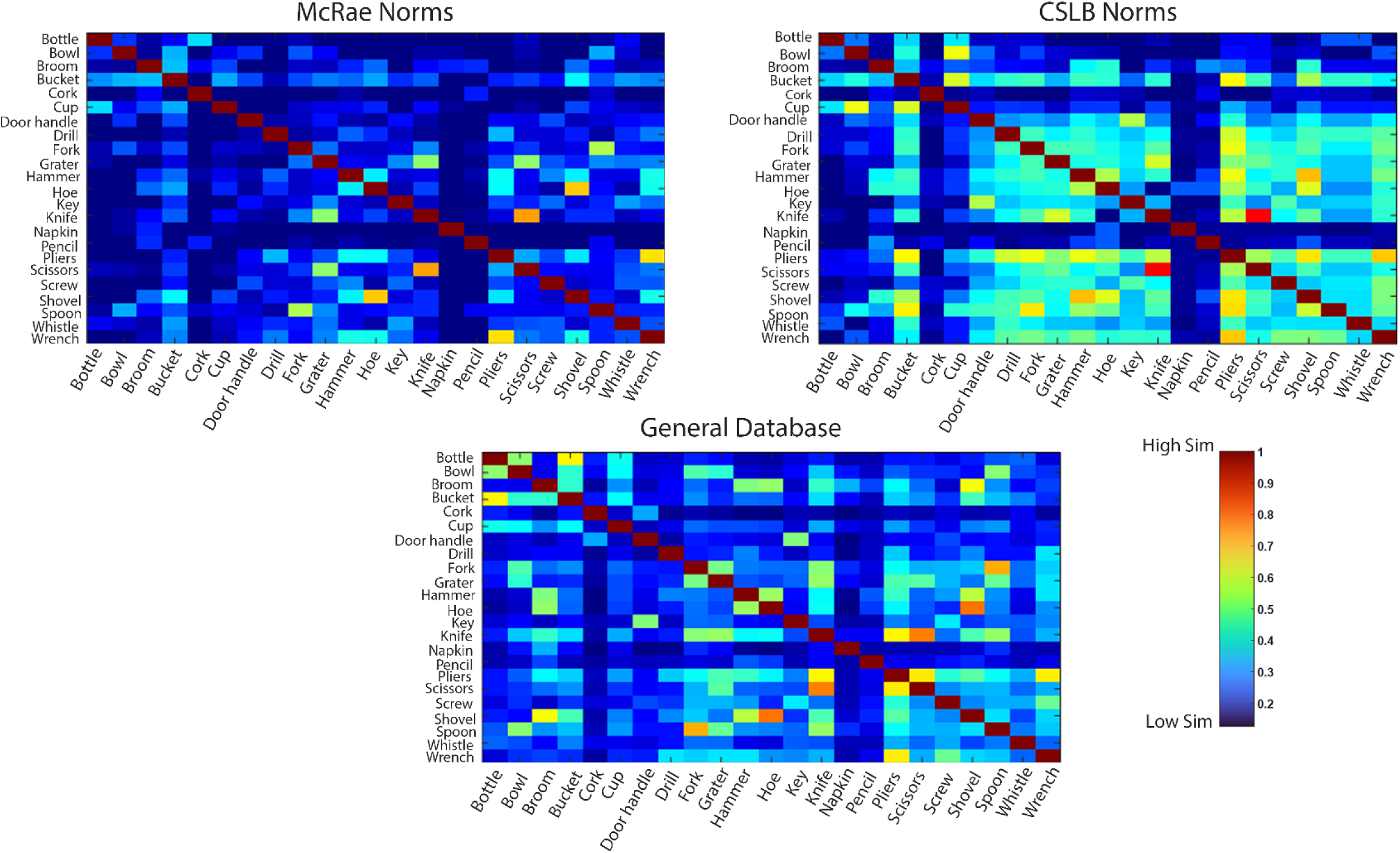
Similarity matrices for 23 manipulable objects appearing in McRae, CSLB, and the General and Action Databases. Rows and columns of the similarity matrices are ordered alphabetically. Colors represent pairwise cosine similarity, which varies between 0 (Low Similarity) and 1 (High Similarity). Note: In CSLB norms, “wrench” is named “spanner”.

### Differences in the Feature Types

Having a database specific to a particular category allowed us to study and disentangle the set of properties that are important to that category (in this case, manipulable objects), which serves as a proxy for understanding fine-grained conceptual processing. To accomplish this, we divided the features into 4 major different types: 1) visual features, which include form, texture/material, and color, 2) functional features, 3) encyclopedic, and 4) action features (i.e., those presented in the action database). The first three feature types were the most frequent in the general database. We defined feature types following the definitions used by McRae and collaborators (McRae et al., 2005). We then calculated the cosine similarity of the 23 manipulable objects for each of the 4 feature types individually and thus created 4 different similarity matrices. We chose the 23 objects that were common between our databases and the McRae and CLSB norms to illustrate this approach. After organizing the objects in alphabetical order, we calculated Pearson correlation values between the four matrices. We also used a complete-linkage hierarchical clustering method to organize the objects belonging to different feature types (see Fig. 3).

**Figure 3:**
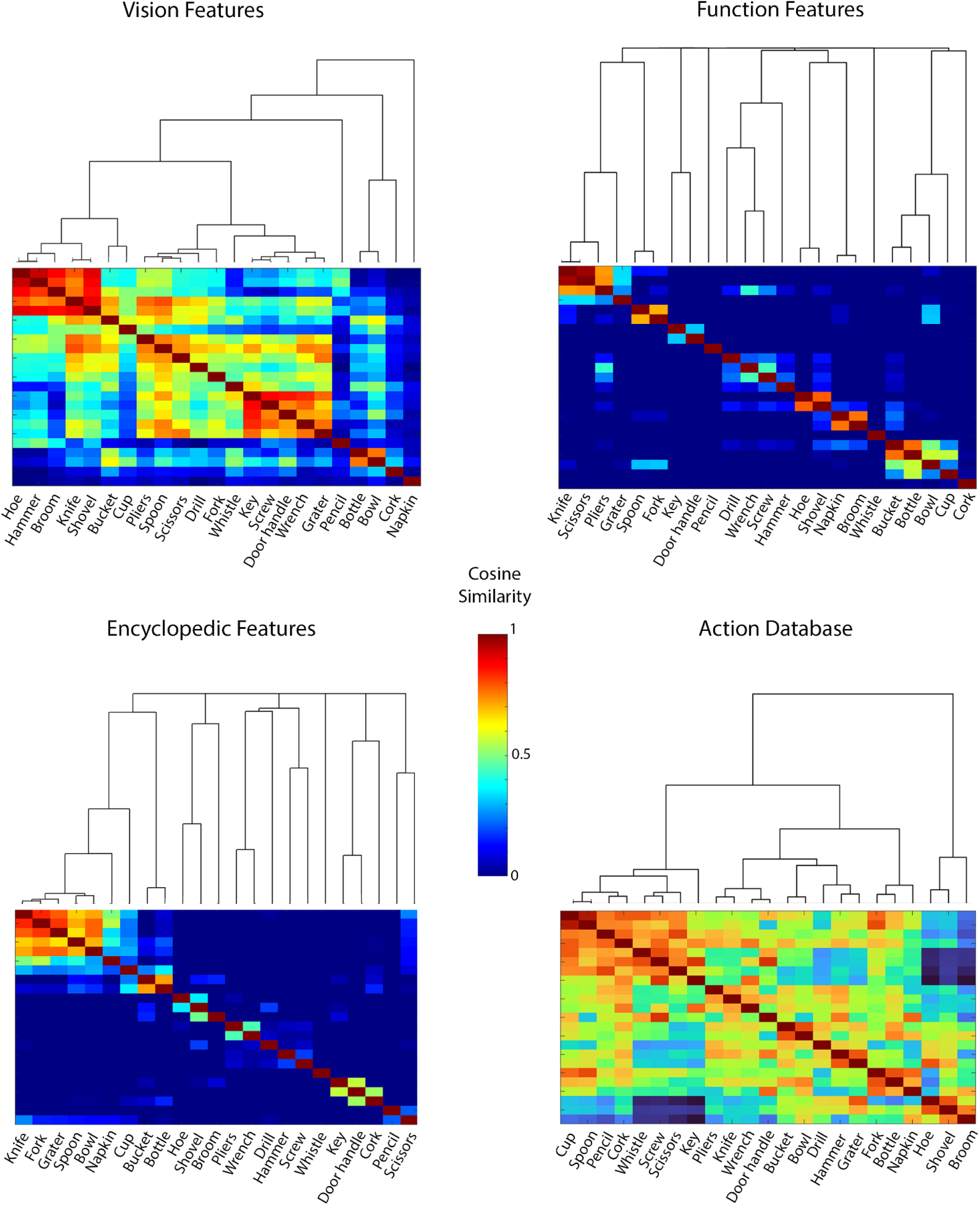
Vision, Function, Encylopedic, and Action. Similarity matrices for the 23 small manipulable objects that were common in the databases. Rows and columns of the similarity matrices are ordered by a complete-linkage hierarchical clustering approach. The colors represent the pairwise cosine similarity, varying from 0 (Low Similarity) to 1 (High Similarity).

## Results

### Norms

We followed the systematic method of McRae and collaborators (McRae et al., 2005) to collate features generated by the participants into lists in both the General and the Action databases. Participants listed the features in Portuguese – both the original Portuguese features and their English translations are presented in our databases. We also present production frequency – i.e., the number of times that a particular feature was mentioned for each object. It is important to note that features with the same meaning were grouped together, even when they were described differently but conveyed the same idea (for a similar method see McRae et al., 2005). To group these features, we used a Portuguese dictionary and the agreement of two researchers (D.V. and J.A.). We removed expressions such as “can be”, “usually”, “many people”, and “generally” from the features. We also split our features based on the different information that they contained, for instance: “used for transporting liquids” was divided into “used for transporting liquids”, “used for transporting”, and “used for liquids”. When a feature contained two different ideas connected by a conjunction or a disjunction, for instance “is black and yellow”, those features were divided into: “is black” and “is yellow”. There were a few cases where participants mentioned the same feature twice. Occasionally, a few incorrect features were listed in our databases (e.g., participants listed “a movie” to describe a nutcracker). In addition, only features that mentioned by at least 5% of participants were included, which is 7 participants for the General database and 2 participants for the Action database.

In the General database, we classified features based on seven types of categories: taxonomic (e.g., “a kitchen object”), encyclopedic (e.g., “used by teachers” or “found in kitchens”), action (“used by clicking”), function (e.g., “used for eating”), visual form and visual texture/material (e.g., “made of metal” or “has two handles”), visual color (e.g., “is black”), tactile (e.g., “is heavy”, or “is hard”), and sound (e.g., “is loud”). We followed the same classification as McRae et al.’s norms (2005), but we added the category of “action” when features are related to the movements of manipulating an object. Additionally, we renamed “visual form and surface” to “visual form and visual texture/material” because we believe it better encompasses the features in this category.

For both the General and Action databases, we calculated the number of concepts per feature (CPF) to count the number of times that each feature was shared among the 80 objects. We also calculated feature distinctiveness in the same way as McRae and collaborators (i.e., 1/CPF; smaller values indicate that the feature is highly shared among objects). Using feature distinctiveness, we determined the mean distinctiveness for each object, which ranges from 0 to 1, with smaller values indicating that the features of that object are highly shared among the objects. Qualitatively, we classified features as distinctive (D) or non-distinctive (ND). A ND feature is shared by at least three objects and helps to identify the group to which the object belongs. A D feature is specific to an object (occurs once or twice in the database) and is important for identifying the object and distinguishing it from similar ones. For example, when trying to infer an object based on a given feature, the ND feature <has blades> is shared by many objects and allows us to identify the object membership (i.e., it is a cutting object), but it is not enough to identify the object. On the other hand, the distinguishing feature <used for cutting paper> is very specific and easily leads to the inference of a pair of scissors. For both databases, we calculated the number of features, number of shared features among objects in our database, number of distinctive features per object, and mean of distinctiveness. We found more shared than distinctive features in both of our databases. Table 1 shows the average values for both databases.

Another variable that was included in the databases was object familiarity. Participants rated how familiar they were with each object on a scale from 1 to 9. In the database, we presented an average and standard deviation of these ratings for each object. The least familiar object was “drill bit” (M [Mean] = 5.1; SD [Standard Deviation] = 2.2), while the most familiar object was “glass” (M = 8.9; SD = 0.3).

Finally, we measured the similarity between two objects based on the shared features and their production frequency. To compute pairwise similarity matrices for both databases, we calculated the cosine similarity using 80 objects x 807 unique features for the General database and 80 objects x 100 unique features for the Action database. We computed the cosine similarity by taking the dot product between two object vectors and dividing it by the product of their lengths. The resulting values range from 0 to 1, with values closer to 1 indicating that two objects have smaller angles and are more similar to each other. These matrices are available along with the databases (a subset of 23 manipulable objects is represented in Figure 2 – General database, and Figure 3 – Action database).

The General and Action databases, the statistics for each manipulable object, and the cosine similarity between objects for each database are available upon request. To obtain these materials, please send an email to the corresponding author.

### Comparison with McRae and CSLB norms

In this section, we report the results from the analysis of the 23 manipulable objects that were in common between our databases and those of McRae and CSLB.

One aspect over which we compare these databases is the overall number of features that were produced per target stimulus. Table 2 shows that our databases have an average of 28.5 features per object in the General database and 12.8 features per object in the Action database. When considering only the General database, there is a statistically significant difference in the number of features obtained when compared with the number presented within the McRae norms (13.7; *t*(44) = –12.6, p < 0.001), but not within the CSLB norms (29.5; *t*(44) = 0.635, *p* = 0.53). Importantly, our General database was compiled from the input of 130 participants, each of whom provided features for all 80 objects, whereas the McRae and CSLB norms were based on the input of 30 participants, each of whom provided features for a subset of concepts. Additionally, in the CSLB norms, the authors guided participants in the features that they should mention.

**Table 2:** Mean and standard deviation (SD) of the Number of features per object, Number of shared features, Number of distinctive features and their percentages for McRae and CSLB norms, and General and Action databases.

|  | Sample | No of features | No of shared features | Shared features % | No of distinctive features | Distinctive features % |
| --- | --- | --- | --- | --- | --- | --- |
| McRae | 30 | 13.7 (3.5) | 4.6 (2.1) | 33.5% | 9.1 (3.5) | 66.5% |
| CSLB | 30 | 29.5 (6.4) | 13.4 (4.3) | 45.5% | 16.1 (7.5) | 54.5% |
| General Database | 130 | 28.5 (4.2) | 15.2 (4.1) | 53.4% | 13.3 (3.2) | 46.6% |
| Action Database | 32 | 12.8 (1.8) | 11 (1.8) | 86.1% | 1.78 (1.4) | 13.9% |

Another important aspect relates to the number of features that are shared or distinctive within these databases. In the General database, 53.4% of the features are shared (i.e., occurring in three concepts or more) and 46.6% are distinctive. In the Action database, 86.1% of features are shared and only 13.9% are distinctive. These values are quite different from McRae norms that presented more distinctive than shared features (i.e., 33.5% are shared and 66.5% distinctive). In contrast, the CSLB norms are more similar to the General database – 45.5% of features are shared and 54.5% are distinctive.

Despite the differences between our General Database and the McRae and CSLB norms, the General database shows a significant and high correlation with both the McRae (*r* = 0.9, p < 0.001) and CSLB (*r* = 0.8, p < 0.001) norms, as measured by object similarity within each database (on the 23 objects that are in common between the databases). The strong correlations can be seen in the high level of similarity demonstrated in Figure 2. We also conducted hierarchical clustering analysis between the 23 manipulable objects in the three databases (See Fig. S1). In the General database, similar objects were clustered together indicating a clear relationship with semantic similarity. For instance, objects that are used as containers were grouped together (i.e., bucket, bottle, bowl and cup). The same can be observed for objects that are used for cutting (i.e., grater, pliers, knife and scissors), or objects found in garages (i.e., drill, hoe, shovel, hammer, and broom). Overall, our databases are interpretable in terms of semantic similarity, and the interpretability of the General database is comparable to that observed for the McRae and CLSB norms (See Fig. S1).

Furthermore, we conducted an analysis to compare the number and percentage of features for each feature type across the 23 manipulable objects that were common between databases (See Table 3). The three most common types of features in the General database were visual, functional, and encyclopaedic, which is similar to what is observed in the McRae and CSLB norms. However, unlike the McRae and CSLB norms, the General database had a more even distribution of features across different types. The General database had a lower percentage of functional features compared to other norms, and the difference between the number of functional and visual features was larger. Moreover, this analysis revealed that action features were present only in our databases and not on the published databases. Overall, the larger number of visual, functional, encyclopaedic, and action features in our databases allows us to individually study and disentangle these four types of features.

**Table 3:** Percentage and Number of features (mean and standard deviation in brackets) per each type of feature for the 23 small manipulable objects.

|  |  | Taxonomic | Encyclopedic | Function | Visual<br>Perceptual* | Other<br>Perceptual# | Action |
| --- | --- | --- | --- | --- | --- | --- | --- |
| McRae | Percentage | 5.8% | 12.4% | 35.8% | 41.6% | 5.1% | — |
|  | No of features | 0.8 (0.7) | 1.7 (1.1) | 4.9 (1.8) | 5.7 (1.9) | 0.7 (1) | — |
| CSLB | Percentage | 5.8% | 16.9% | 30.5% | 39.3% | 7.8% | — |
|  | No of features | 1.7 (1.2) | 5 (2.3) | 9 (3.2) | 11.6 (3.1) | 2.3 (1.7) | — |
| General<br>Database | Percentage | 7% | 23.2% | 21.8% | 37.2% | 5.3% | 5.6% |
|  | No of features | 2 (1.3) | 6.6 (2.3) | 6.2 (2.1) | 10.6 (2.8) | 1.5 (0.8) | 1.6 (1) |
| Action<br>Database | Percentage | — | — | — | — | — | 100% |
|  | No of features | — | — | — | — | — | 12.8 (1.8) |

\* Visual Perceptual is the combination between Visual form and surface (equivalent to our Visual form and visual texture/material) and Visual color in the McRae and General Databases. # Other Perceptual is a combination between Tactile and Sound in the McRae and General Databases.

### Differences in the Feature Types

We analyzed the four types of features and found weak to moderate correlations among them. Specifically, vision and encyclopedic features (r = 0.41, p < 0.001), vision and action features (r = 0.24, p < 0.001), function and action features (r = 0.47, p < 0.001), action and encyclopedic features (r = 0.46, p < 0.001), and vision and function features (r = 0.51, p < 0.001) are all significantly and moderately correlated. In contrast, function and encyclopedic features were strongly correlated (r = 0.75, p < 0.001). Moreover, the clustering solution provided a clear and highly interpretable set of clusters that are specific to each feature type. For vision features, objects were clustered together dependent on whether they were elongated with a wooden handle (i.e., hoe, hammer, broom, shovel, and knife), made of metal and/or plastic (i.e., scissors, pliers, spoon, drill, and fork), and entirely made of metal (i.e., key, screw, door handle and wrench). For function features, objects used for cutting (knife, scissors, pliers, and grater), transporting liquids (bucket, bottle, bowl, and cup), repairing (drill, wrench, screw, hammer) and cleaning (napkin and broom) were clustered together. For encyclopedic features, clustering was based on the shared context between objects. For example, objects were organized based on the kitchen context (knife, fork, grater, spoon, bowl, napkin, cup, bucket, and bottle), the school/office context (pencil and scissors), and garage objects typically found in toolboxes (pliers, wrench, drill, hammer, and screw). For the Action database, there are two large clusters representing objects used with one hand (the majority) or two hands (e.g., hoe, shovel, and broom). Among objects used with one hand, the cluster solution further divides objects into those requiring precise movements (e.g., cup, whistle, or scissors) and those requiring full-handed grasp (e.g., knife, door handle, or hammer). These results suggest that these four types of features contain different information, as the object matrices were distinct and meaningful (See Fig. 3).

## Discussion

Here, we present two databases that focus on 80 familiar manipulable objects: 1) the General database – in which participants freely generated features; and 2) the Action database – where participants generated features related to the manner in which these objects are used. We created an Action database because information about object manipulation has been shown to be important for our understanding of manipulable objects (e.g., Almeida et al., 2018; Almeida, Mahon, & Caramazza, 2010; Almeida, Mahon, Nakayama, & Caramazza, 2008; Almeida et al., 2014; Buxbaum & Saffran, 2002; Garcea et al., 2013; Valério et al., 2021). In order to collate features from the many participants into the norms presented here, we followed the systematic method created by McRae and collaborators (2005), and only included features mentioned by at least 5% of our sample. Our features cover many aspects of object knowledge, including action features –an advantage over other feature databases. Across our two databases, the most common feature types, ordered by frequency, were visual, action, encyclopedic and functional, with enough features to study each type individually. In the General database, we have a lower percentage of functional features compared to other norms, which was likely due to the even distribution of features across feature types. This distribution might be a result of the nature of our task, where participants generated within-category features, and consequently they had a tendency to produce more general features important to the concept and less focused on the differences between categories. In a feature generation task between categories, such as the one used by McRae and collaborators for instance, participants tended to generate more functional features to describe manipulable objects, perhaps because participants were more focused on the features that distinguish this category from others, such as animals or fruits and vegetables. Analogously, if participants were presented with many manipulable objects and fruits, they would most likely describe fruits with more taste-related features, because taste may be one of feature types that distinguishes between fruits and small manipulable objects.

Additionally, we found our databases to contain more shared features than distinctive ones. Interestingly, according to the literature, manmade objects are typically considered to be very different from each other (e.g., when compared to animals), and present more distinctive features (Cree & McRae, 2003; Tyler, Moss, Durrant-Peatfield, & Levy, 2000). This raises the question of whether these previous results relate to the fact that manmade objects do have more distinctive features, or, alternatively, whether those data were obtained in the context of stimuli from other categories. Our data clearly suggest the latter to be the case. Importantly, many semantic memory theories are based on neuropsychological data from category-specific deficits presented by individuals with brain damage, who demonstrate difficulties in performing within-category distinctions (Capitani et al., 2003; Caramazza & Mahon, 2003).Therefore, this suggests that we should at least also focus on within-category processing in order to fully understand the type of object knowledge we store and use when processing objects.

Interestingly, the large number of shared features (53.1% in the General database and 86.1% in the Action database) allows us to study the similarity between concepts or specific features that are common to many objects, such as “made of metal” or “precise movements/use of fingers”. On the other hand, the presence still of a significant number of distinctive features, particularly in the General database, allows us to identify which features are important for discriminating between similar objects.

Finally, we compared the 23 manipulable objects that are common to both the General database and the McRae and CSLB norms. We found a high correlation between the databases (Fig. 2), and confirmed that the General database reliably codes for object similarity in the same fashion as the established databases in the literature (See Fig. S1 of the Supplementary Material). Moreover, for these 23 objects, we analyzed the differences between the three most frequent types of features in the General database: vision, function and encyclopedic, as well as the action features from the Action database. Our results showed that objects were grouped differently depending on the type of feature. For example, based on similarity computed over vision features, a knife was grouped with elongated objects that have a wooden handle (i.e., hoe, hammer, broom, shovel); based on function features, a knife was grouped with cutting objects (i.e., scissors, pliers and a grater); using encyclopedic features, a knife was grouped with kitchen objects (i.e., a fork, grater, spoon, and bowl), and finally, based on action features, a knife was clustered with objects used with one-hand and palmar grasp (i.e., pliers, wrench, and door handle; See Fig. 3). Interestingly, we found a closer relation between function and encyclopedic features, likely because objects that have similar functions (e.g., used for writing or preparing meals) are also used in similar contexts (e.g., at school or in the kitchen). These results and differences may be important for future studies relating to how these different types of information (e.g., action, function, and vision) are processed in the service of manipulable object perception (and object perception in general) both in patient (Buxbaum & Saffran, 2002; Buxbaum, Veramontil, & Schwartz, 2000; Goodale et al., 1994; for a review see: Mahon & Caramazza, 2003), and in healthy populations (Chen, Garcea, & Mahon, 2016; Lesourd et al., 2023; Peelen & Caramazza, 2012). Therefore, by investigating these differences between types of features, we may be able to further disentangle the organization of object knowledge in the brain.

### Experiment 2: Feature Verification Task

In Experiment 2, our goal was to explore whether a different set of participants could recognize and associate the previously generated features with specific manipulable objects, and thus test the reliability of our feature database. To do so, we manipulated feature production frequency values, as previous studies have shown that this measure reflects the importance of a feature for a particular concept (Ashcraft, 1976; McRae, De Sa, & Seidenberg, 1997). We expected that object-specific highfrequency features are more salient, and thus participants will be faster and more accurate to associate these features with the target object. We measured reaction times (RTs) and recorded the accuracy of feature verification performance for features with high and low frequency production in a feature verification task. In this task, participants were asked to determine whether a given feature was true (or false) of a specific object. We expected that participants would show similar performances for both the General and the Action databases.

To accomplish these goals, we performed three tasks: 1) test the effect of production frequency in a feature verification task for the General database in a laboratory environment; 2) replicate task 1 with a new group of online participants; and 3) test the effect of production frequency in a feature verification task with yet another group of participants online using the Action database.

## Method

### Participants

Thirty volunteers, 25 women, performed the feature verification task using features from the General database in a laboratory environment. All but three participants were right-handed. Their average age was 19.4 years (ranging from 18 to 25). Twenty-one individuals, 19 of whom were women and all of whom were right-handed, participated in the online feature verification task with the General database. Their average age was 19.8 (ranging from 18 to 31). In the Action Database Online condition, 30 volunteers participated, 26 of whom were women, 26 of whom were right-handed, and one of whom was ambidextrous. Their average age was 21.5 (ranging from 18 to 49).

All participants were European Portuguese native speakers. All participants were naïve to the goal of the experiment, were undergraduate psychology students, and received a course bonus for their participation in this experiment. We obtained written informed consent from all participants according to the Ethics committee of the Faculty of Psychology and Educational Sciences, University of Coimbra.

### Procedures

In the three conditions of Experiment 2, participants saw <object-feature> pairs and were asked to determine whether the feature was true or false for that object, while we measured RTs and accuracy. We presented features with high or low frequency, based on how frequently each feature was mentioned in our databases. In this experiment, the high-frequency features belonged to the 75% most frequent features of each object, while low-frequency features represented the 25% least frequent features. We controlled the number of characters between conditions.

In each trial, an <object-feature> pair appeared on the screen, both stimuli presented in lowercase letters. The different conditions were presented in a random order. The trial began with a fixation cross that remained on the screen for 0.5 seconds. Then, the object appeared 0.350 seconds before the feature (the stimulus onset asynchrony). The object name remained on the screen throughout the trial. Participants responded using a keyboard, with assignments counterbalanced across participants (i.e., half of the participants pressed “p” for true pairs and “q” for false pairs, and vice-versa). The monitor refresh rate was 60 Hz.

For the General database, each participant completed two sessions on different days, each lasting approximately 40 minutes, resulting in a total of 1280 trials per participant. During each session, every object appeared 8 times, paired with different features – 4 as true and 4 as false pairs. For the true pairs, we selected two of the features per object that were high-frequency features and two that were low-frequency. For the false pairs, we used the same features as in the true pairs, but paired them with an object they did not relate with. To illustrate, consider the example of scissors: the features “used for cutting” and “made of plastic” were high-frequency, whereas “a school object” and “different sizes” were low-frequency. The false features were “made of paper”, “associated with wind”, “associated with cars”, and “used for weighing on a balance scale”. For the second session, we used different features. See Figure 1B for three trial examples of high and low-frequency features (i.e., true conditions) and false features.

In the Action Database Online, participants completed only one session, which took approximately 40 minutes. Each object appeared eight times in a session. Using the scissors example again, the features “precise movements/use of fingers” and “opening and closing movements” were high-frequency, whereas “use of three fingers” and “movement of approaching and moving away from the fingers” were low-frequency. The false features were “palmar_grasp/full-handed_grasp”, “moves upwards”, “slow movements”, and “several movements”. Each <object-feature> pair appeared only once during the experiment.

For the in-person lab experiment, we used MATLAB and Psychtoolbox (Kleiner et al., 2007) to control for stimuli presentation and obtain behavioral data, whereas for the online experiments we used OpenSesame (Mathôt, Schreij, & Theeuwes, 2012).

### Analysis

RTs that were below 0.150 seconds (i.e., too fast) or above 6 seconds were discarded and considered as incorrect responses. To analyze the RTs and accuracy, we used two separate within-subject one-way ANOVAs with frequency type (high vs low frequency) as factor. Paired t-tests were also performed. We only analyzed the true <object-feature> pairs.

## Results

### For the three tasks, we analyzed RTs and percent accuracy independently

In the General database Lab-environment task, there was a statistical difference between the high and low production frequency features when judging whether a feature belongs to an object. Reaction times were significantly slower for low-frequency features compared to high-frequency features (*F*(1,29) = 131.03, *P* < 0.0001, and partial η² = 0.82; *t*(29) = –11.47, *P* < 0.0001; see Table 4 for RTs). Participants were also less accurate for low-frequency features compared to high-frequency features (*F*(1,29) = 173.87, *P* < 0.0001, and partial η² = 0.86; *t*(29) = –13.19, *P* < 0.0001; see Table 4 for accuracy scores).

**Table 4:** Results of Experiment 2, for Reaction Times (RTs) and the Percentage of Errors for the true pairs.

|  |  | General Database Lab |  | General Database Online |  | Action Database Online |  |
| --- | --- | --- | --- | --- | --- | --- | --- |
|  |  | High Freq. | Low Freq. | High Freq. | Low Freq. | High Freq. | Low Freq. |
| RTs (sec.) | Mean | 1.18 | 1.33 | 1.16 | 1.29 | 2.05 | 2.30 |
|  | SD | 0.27 | 0.29 | 0.17 | 0.20 | 0.47 | 0.60 |
|  | SEM | 0.05 | 0.05 | 0.04 | 0.04 | 0.09 | 0.11 |
|  | Max. | 1.91 | 2.06 | 1.46 | 1.54 | 3.24 | 4.02 |
|  | Min | 0.75 | 0.86 | 0.82 | 0.92 | 0.98 | 1.12 |
| Errors (%) | Mean | 10.78 | 21.21 | 17.91 | 34.91 | 24.77 | 37.92 |
|  | SD | 4.84 | 7.18 | 7.48 | 9.09 | 8.27 | 8.79 |
|  | SEM | 0.88 | 1.31 | 1.63 | 1.98 | 1.51 | 1.87 |
|  | Max. | 22.19 | 36.56 | 35.00 | 63.44 | 49.38 | 62.50 |
|  | Min. | 3.75 | 7.19 | 7.19 | 23.13 | 14.38 | 19.38 |

We replicated these results in the General Database Online task, where we again show a statistical difference between high-and low-frequency features for both reaction times (*F*(1,20) = 120.58, *P* < 0.0001, and partial η²= 0.86; *t*(20) = –10.98, *P* < 0.0001) and percent accuracy (*F*(1,20) = 256.53, *P* < 0.0001, and partial η²= 0.93; *t*(20) = –16.02, *P* < 0.0001; see Table 4).

In the Action database Online, we found that reaction times (*F*(1,29) = 37.65, *P* < 0.0001, and partial η² = 0.57) and accuracy (*F*(1,29) = 118.30, *P* < 0.0001, and partial η²= 0.80) differed significantly for the frequency feature. Participants were statistically faster (*t*(29) = –6.14, *P* < 0.0001) and more accurate (*t*(29) = –10.88, *P* < 0.0001) when verifying high-frequency features related to the manner of manipulating objects compared to low-frequency features (see Fig. 4 and Table 4).

**Figure 4:**
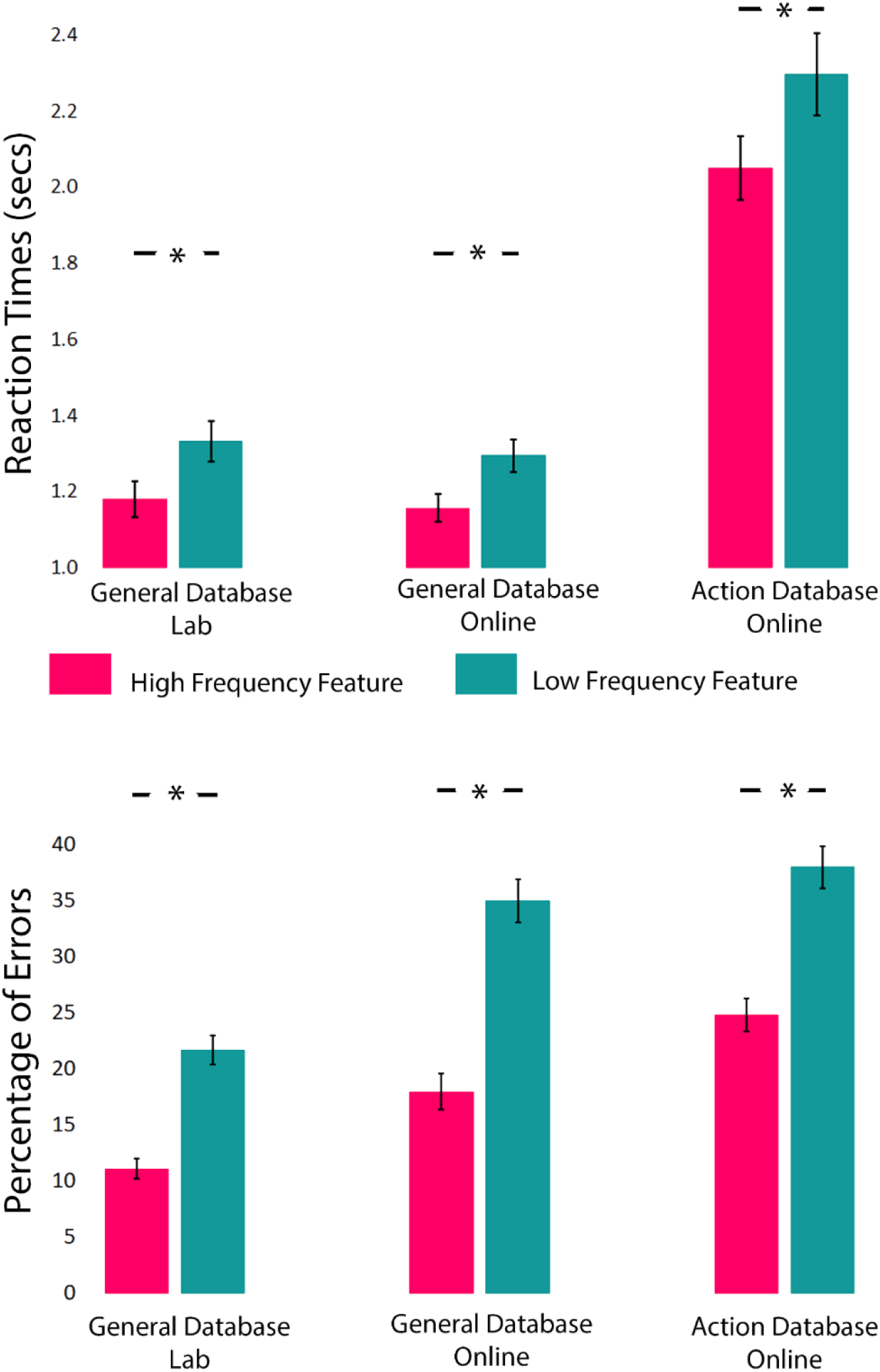
Reaction Times and Percentage of Errors for the feature verification task. Here we show RTs and percent accuracy performance in the three tasks as a function of the production frequency of the features used. * Indicates statistical significance at *P* < 0.0001

## Discussion

Across the three feature verification tasks, we found that participants were faster and more accurate with high-frequency features than low-frequency features (Table 4, and Fig.4). These findings are in line with previous studies (Ashcraft, 1976; McRae et al., 1997) and are consistent across conditions and participants (See Fig. S3 of the Supplementary Material).

There are, however, differences between our results and those in the literature, namely in what concerns the percentage of errors, as our experiments show a relatively high number of errors. This is especially true for the action task. This may be so because verifying these kinds of features is a complicated task (as indexed by the higher RTs in this experiment). Moreover, another possible reason for the high percentage of errors in our feature verification task is that we use only manipulable objects – i.e., people are doing within category decisions – many of them with high similarity. This contrasts with most tasks reported in the literature, which present multiple concepts from different categories. For instance, McRae and collaborators (1997) used different categories (e.g., animals, vehicles, small manipulable objects, fruits and vegetables) and they found an average of errors of 4.3% and 13% of “strong” and “weak” production frequency, respectively. Therefore, the greater difficulty of our task is most likely the reason for this discrepancy in percentage of errors. Importantly, we found no differences in reaction time between experiments performed in the lab environment and at home (General database).

Most importantly, these experiments are a demonstration that our feature database is on par with the main databases in the field in its ability to explain decisions related to object knowledge.

### General Discussion

Semantic norms have played a key role in testing and developing theories of semantic representation and object recognition. In recent years, research has focused on how to differentiate living and non-living things in semantic memory, or to distinguish between different semantic categories namely animals, manipulable objects, body parts, fruits and vegetables among others. The field has, however, given much less attention to within-category differences and processing. Importantly, within-category performance is, in fact, what is affected in the kinds of category-specific deficits that have been used to inform theories of semantic memory. Our goal here was to study how fine-grained object specific content is represented and processed in semantic memory. To do this, we focused on within-category feature norms as a way to strip out general and gross categorical distinctions and focus on hard and more subtle distinctions between objects.

We used the category of manipulable objects as a case study, and asked two samples of participants to generate features related to 80 familiar objects that are frequently used in daily life. As result, we compiled, on average, 27.5 and 12.9 features per object in the General and Action databases, respectively. This provides a comprehensive list of features that allows us to describe various aspects of an object, such as features related to visual appearance (i.e., form and surface information), color, function, manner of use, encyclopedic knowledge (i.e., context and associations), and other sensory information (i.e., sound and tactile). We differentiate between shared and distinctive features, and because we are using a single category and many objects have close relatives, we found that a considerable number of features are shared among the objects. In fact, the General database contains an almost equal percentage of shared and distinctive features. These results are inconsistent with the McRae norms exhibit a higher number of distinctive features for non-living objects (See Table 2). Indeed, in the literature, non-living objects are frequently thought of as having more distinctive than shared features, while animals display the opposite effect. On the 99 concepts that participants defined as animals in the McRae norms, 81% of their features were shared (e.g., “has four legs”, “has a tail”, or “eats”), and just 7% of the features were functional (e.g., “eaten as pork”, “used for protection”, or “produces milk”). Several factors might explain this effect: 1) the McRae norms have fewer features per concept, with those features being the most frequently used to describe a concept (mentioned by at least 5 out of 30 participants); 2) the majority of these animals were mammals, making them very similar in their appearance – if the database included a more diverse array of animals (such as insects, reptiles, fish and birds) this percentage would likely decrease; 3) functional features were more commonly found in animals that are typically consumed as food (e.g., pigs, cows or chickens) or animals that participants considered as pets (e.g., dog or horse). Thus, if we only considered those animals, the differences between visual and functional would likely vanish; and 4) when participants are required to describe features of many concepts from many different categories, they tended to focus more on the features that distinguish different categories (e.g., visual features that are in common between animals, or what objects are used for when thinking about manipulable objects). Importantly, our results do not support the contention that manipulable object have more distinctive features than shared features, not more functional than visual features. Rather, previous effects suggesting these differences were due to the context of the task used, whereby items from many different categories accentuated putative spurious categorical differences.

We also calculated the similarity between concepts using features from our databases and from the McRae and CSLB norms, and found that the General database is highly correlated and comparable in terms of semantic similarity and interpretability with those norms (See Fig. 2 and Fig. S2 of the Supplementary Material). However, the McRae norms have a smaller number of visual, functional and encyclopedic features per concept, making it difficult to study each type of feature individually. The CSLB norms have a higher number of features per feature type but do not include action features (See Table 3). Therefore, one of the advantages of our databases is the possibility to study visual, functional, encyclopedic and action features individually. Importantly, these kinds of features and the conceptual knowledge associated have been shown to rely on different brain areas (Boronat et al., 2005; Chen et al., 2016; Culham et al., 2003; Lesourd et al., 2023; Peelen & Caramazza, 2012), and are doubly dissociated behaviorally (Garcea & Mahon, 2012; Myung, Blumstein, & Sedivy, 2006) and in patients (Buxbaum & Saffran, 2002; Garcea et al., 2013; Goodale et al., 1994; Myung et al., 2010). Analysis of these four feature types in terms of object similarity provided with meaningful associations between objects. For instance, in the action database, objects used in a similar way were grouped together, allowing us to differentiate objects used with one or two hands, and those requiring a palmar grasp (i.e., full-handed) or precise movements (i.e., use of fingers).

We also tested the validity of our databases by conducting a feature verification experiment, showing that a different set of subjects can recognize and associate the features present in both databases to particular objects. Specifically, we asked three groups of participants to verify if a feature was true of a particular object, while manipulating production frequency. We found that participants were faster and more accurate with high-frequency features (i.e., those with greater consensus) presented from the General and Action databases. This is likely because high-frequency features are more salient to a particular object and contribute more to its core meaning.

To summarize, most theories of semantic memory focus on the differences between categories to explain how information is processed and represented in the brain. However, little attention has been given to how information is organized within-category. This is particularly problematic because indeed, patients with neurological impairments demonstrate difficulties in performing within-category distinctions. Our databases have the potential to support research on how content-specific information of a single category – manipulable objects – is represented and processed in the brain. This study could also contribute to a better understanding of how the brain efficiently identifies and differentiates among objects, despite significant variations in shapes, colors, materials, and viewpoints. Additionally, it could help disentangle how different types of features are represented within the manipulable objects category.

## Acknowledgements

We would like to thank Ema Leitão, Catarina Senra, and Adriana Martins for their help in data collection and Arthur Pilacinski for his help with OpenSesame.

## Supplementary Material for Semantic feature production norms for manipulable objects

Supplementary Table S1: The 80 manipulable objects used in the General and Action databases

Supplementary Figure S1: Similarity matrices for 23 manipulable objects appearing in McRae, CSLB, and the General and Action Databases.

Supplementary Figure S2: Reaction Times of participants from Experiment 2 in A) General database lab; B) General database online; and C) Action database online.

**Table S1:** The 80 manipulable objects used in the General and Action databases

| English | English | English | English |
| --- | --- | --- | --- |
| 1 – Basketball | 21 – Flashlight | 41 - Mop | 61 - Rubber stamp |
| 2- Board eraser | 22 – Fork | 42 – Nail | 62 - Scissors |
| 3 - Bottle | 22 – Glass | 43 - Nail clipper | 63 - Screw |
| 4- Bottle Cap | 23 -Golf club | 44 - Nail file | 64 - Shaver |
| 5 – Bottle opener | 24 - Grater | 45 - Napkin | 65 - Shopping cart |
| 6 - Bowl | 25 -Hairbrush | 46 - Needle | 66 - Shovel |
| 7 - Broom | 26 - Hammer | 47 - Nutcracker | 67 - Spinning top |
| 8 – Bucket | 27 - Hand Blender | 48 - Paddle | 68 - Sponge |
| 9 – Candle holder | 28 - Hand Fan | 49 – Paintbrush | 69 - Spoon |
| 10 – Citrus squeezer | 29 - Hanger | 50 - Paper Clip | 70 - Spray bottle |
| 11 – Clothespin | 30 - Hoe | 51 - Peeler | 71 - Stapler |
| 12 - Computer mouse | 31 - Hole punch | 52 - Pencil | 72 - Swiss knife |
| 13 - Cork | 32 – Horn | 53 - Pencil eraser | 73 - Syringe |
| 14 - Cup | 33 - Jug | 54 - Pencil sharpener | 74 - Toothbrush |
| 15 - Dart | 35 - Key | 55 - Pepper grinder | 75 - Tweezers |
| 16 - Door handle | 36 - Knife | 56 - Pastry bag | 76 - Weights |
| 17 - Drill | 37 - Lifebuoy | 57 - Pliers | 77 – Whisk |
| 18 - Drill Bit | 38 - Lighter | 58 - Plunger | 78 - Whistle |
| 19 - Dryer | 39 - Magnifying glass | 59 - Racket | 79 - Wooden spoon |
| 20 - Fishing rod | 40 - Match | 60 - Rolling pin | 80 - Wrench |

**Figure S1:**
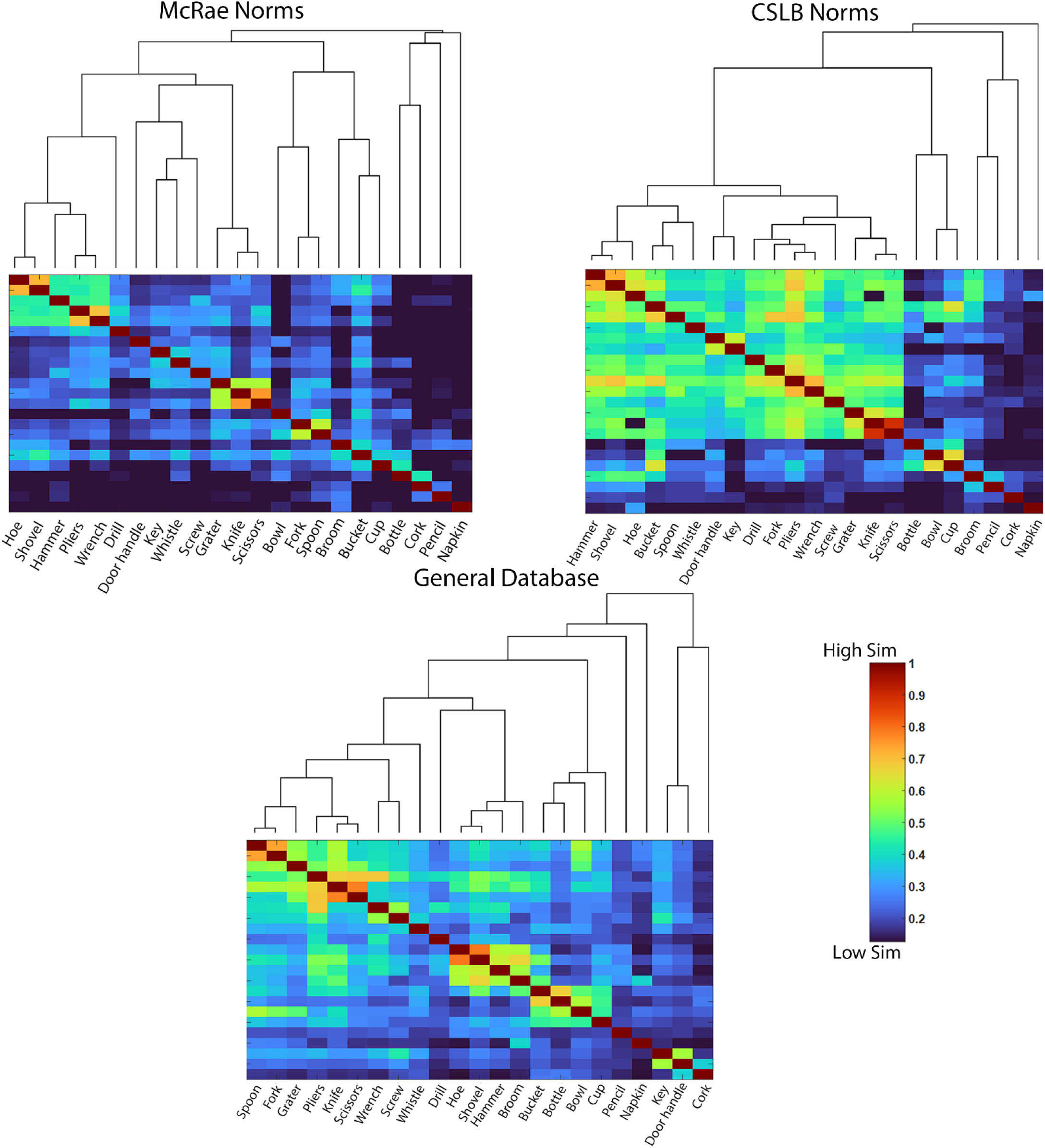
Similarity matrices for 23 manipulable objects appearing in McRae, CSLB, and the General and Action Databases. Rows and columns of the similarity matrices are ordered by a complete-linkage hierarchical clustering. The colors represent the pairwise cosine similarity, varying from 0 (Low Similarity) to 1 (High Similarity).

**Figure S2:**
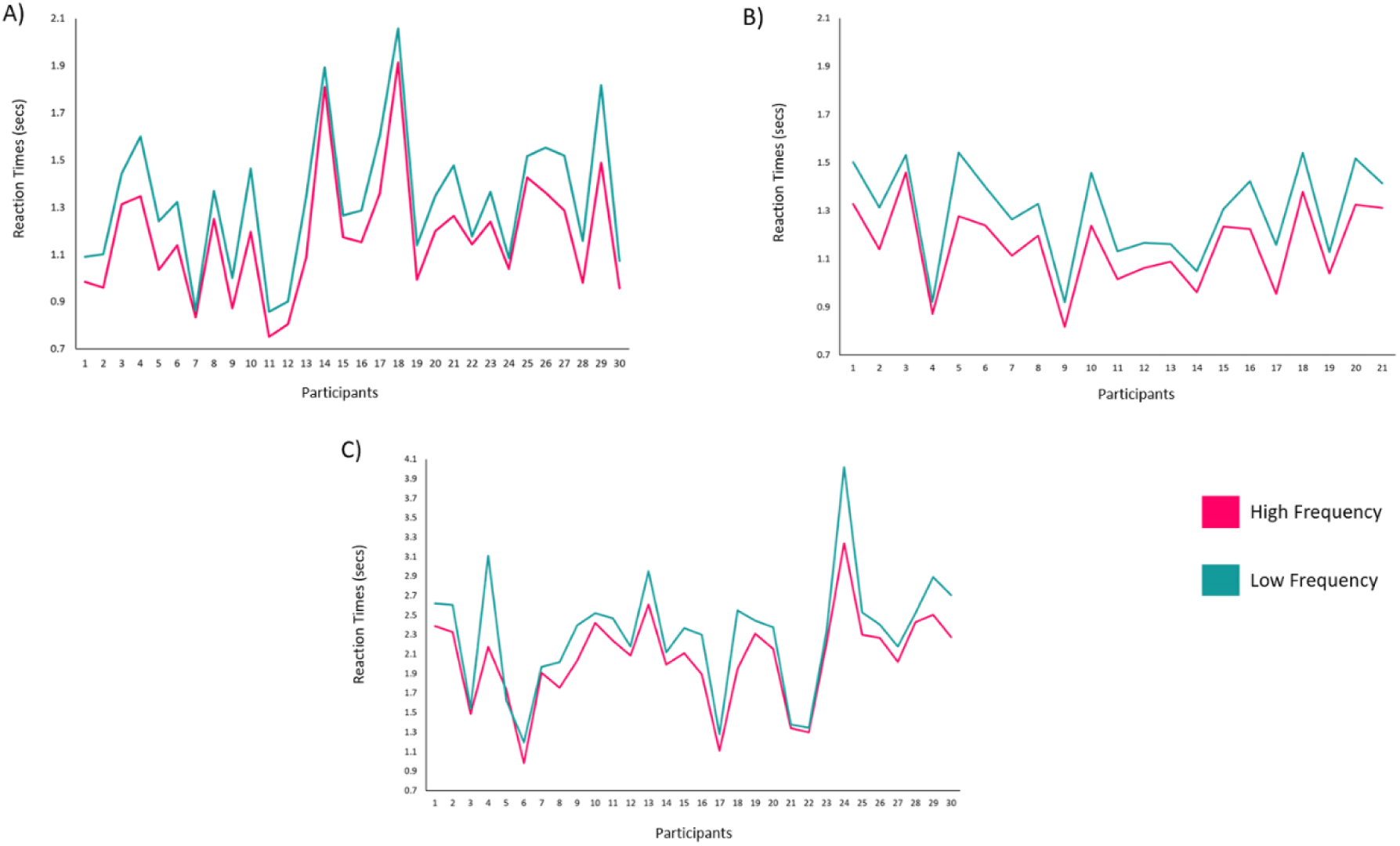
Reaction Times of participants from Experiment 2 in A) General database lab; B) General database online; and C) Action database online. Results for High and Low-frequency true pairs are represented in pink and blue, respectively.

## References

Almeida, J., Amaral, L., Garcea, F. E., Aguiar de Sousa, D., Xu, S., Mahon, B. Z., & Martins, I. P. (2018). Visual and visuomotor processing of hands and tools as a case study of cross talk between the dorsal and ventral streams. Cognitive Neuropsychology, 35(5–6), 1–16. https://doi.org/10.1080/02643294.2018.1463980

Almeida, J., Mahon, B. Z., & Caramazza, A. (2010). The role of the dorsal visual processing stream in tool identification. Psychological Science: A Journal of the American Psychological Society / APS, 21(6), 772–778. https://doi.org/10.1177/0956797610371343

Almeida, J., Mahon, B. Z., Nakayama, K., & Caramazza, A. (2008). Unconscious processing dissociates along categorical lines. Proceedings of the National Academy of Sciences of the United States of America, 105(39), 15214–15218. https://doi.org/10.1073/pnas.0805867105

Almeida, J., Mahon, B. Z., Zapater-Raberov, V., Dziuba, A., Cabaço, T., Marques, J. F., & Caramazza, A. (2014). Grasping with the eyes: The role of elongation in visual recognition of manipulable objects. *Cognitive*, Affective and Behavioral Neuroscience, 14(1), 319–335. https://doi.org/10.3758/s13415-013-0208-0

Ashcraft, M. H. (1976). Priming and property dominance effects in semantic memory. Memory & Cognition, 4(5), 490–500. https://doi.org/10.3758/BF03213209

Boronat, C. B., Buxbaum, L. J., Coslett, H. B., Tang, K., Saffran, E. M., Kimberg, D. Y., & Detre, J. A. (2005). Distinctions between manipulation and function knowledge of objects: Evidence from functional magnetic resonance imaging. Cognitive Brain Research, 23(2–3), 361–373. https://doi.org/10.1016/j.cogbrainres.2004.11.001

Buxbaum, L. J., & Saffran, E. M. (2002). Knowledge of object manipulation and object function: Dissociations in apraxic and nonapraxic subjects. Brain and Language, 82(2), 179–199. https://doi.org/10.1016/S0093-934X(02)00014-7

Buxbaum, L. J., Veramontil, T., & Schwartz, M. F. (2000). Function and manipulation tool knowledge in apraxia: Knowing ‘what for’ but not ‘how.’ Neurocase, 6(2), 83–97. https://doi.org/10.1080/13554790008402763

Capitani, E., Laiacona, M., Mahon, B., & Caramazza, A. (2003). What Are the Facts of Semantic Category-Specific Deficits? a Critical Review of the Clinical Evidence. Cognitive Neuropsychology, 20(3–6), 213–261. https://doi.org/10.1080/02643290244000266

Caramazza, A., & Mahon, B. Z. (2003). The organization of conceptual knowledge: The evidence from category-specific semantic deficits. Trends in Cognitive Sciences, 7(8), 354–361. https://doi.org/10.1016/S1364-6613(03)00159-1

Chen, Q., Garcea, F. E., & Mahon, B. Z. (2016). The Representation of Object-Directed Action and Function Knowledge in the Human Brain. Cerebral Cortex, 26(4), 1609–1618. https://doi.org/10.1093/cercor/bhu328

Clarke, A., Taylor, K. I., Devereux, B., Randall, B., & Tyler, L. K. (2013). From perception to conception: How meaningful objects are processed over time. Cerebral Cortex, 23(1), 187–197. https://doi.org/10.1093/cercor/bhs002

Cree, G. S., & McRae, K. (2003). Analyzing the Factors Underlying the Structure and Computation of the Meaning of Chipmunk, Cherry, Chisel, Cheese, and Cello (and many Other Such Concrete Nouns). Journal of Experimental Psychology: General, 132(2), 163–201. https://doi.org/10.1037/0096-3445.132.2.163

Culham, J. C., Danckert, S. L., DeSouza, J. F. X., Gati, J. S., Menon, R. S., & Goodale, M. A. (2003). Visually guided grasping produces fMRI activation in dorsal but not ventral stream brain areas. Experimental Brain Research, 153(2), 180–189. https://doi.org/10.1007/s00221-003-1591-5

de Deyne, S., Verheyen, S., Ameel, E., Vanpaemel, W., Dry, M. J., Voorspoels, W., & Storms, G. (2008). Exemplar by feature applicability matrices and other Dutch normative data for semantic concepts. Behavior Research Methods, 40(4), 1030–1048. https://doi.org/10.3758/BRM.40.4.1030

Devereux, B. J., Tyler, L. K., Geertzen, J., & Randall, B. (2014). The Centre for Speech, Language and the Brain (CSLB) concept property norms. Behavior Research Methods, 46(4), 1119–1127. https://doi.org/10.3758/s13428-013-0420-4

Duarte, L. R., Marquié, L., Marquié, J. C., Terrier, P., & Ousset, P. J. (2009). Analyzing feature distinctiveness in the processing of living and non-living concepts in Alzheimer’s disease. Brain and Cognition, 71(2), 108–117. https://doi.org/10.1016/j.bandc.2009.04.007

Garcea, F. E., Dombovy, M., & Mahon, B. Z. (2013). Preserved tool knowledge in the context of impaired action knowledge: implications for models of semantic memory. Frontiers in Human Neuroscience, 7(120), 1–18. https://doi.org/10.3389/fnhum.2013.00120

Garcea, F. E., & Mahon, B. Z. (2012). What is in a tool concept? Dissociating manipulation knowledge from function knowledge. Memory and Cognition, 40(8), 1303–1313. https://doi.org/10.3758/s13421-012-0236-y

Garrard, P., Lambon Ralph, M. A., Hodges, J. R., & Patterson, K. (2001). Prototypicality, distinctiveness, and intercorrelation: Analyses of the semantic attributes of living and nonliving concepts. Cognitive Neuropsychology, 18(2), 125–174. https://doi.org/10.1080/02643290125857

Goodale, M. A., Meenan, J. P., Bülthoff, H. H., Nicolle, D. A., Murphy, K. J., & Racicot, C. I. (1994). Separate neural pathways for the visual analysis of object shape in perception and prehension. Current Biology, 4(7), 604–610. https://doi.org/10.1016/S0960-9822(00)00132-9

Hillis, A. E., & Caramazza, A. (1991). Category-specific naming and comprehension impairment: A double dissociation. Brain, 114, 2081–2094. https://doi.org/10.1093/brain/114.5.2081

Kivisaari, S. L., van Vliet, M., Hultén, A., Lindh-Knuutila, T., Faisal, A., & Salmelin, R. (2019). Reconstructing meaning from bits of information. Nature Communications, 10(1), 1–11. https://doi.org/10.1038/s41467-019-08848-0

Kleiner, M., Brainard, D., Pelli, D., Ingling, A., Murray, R., & Broussard, C. (2007). What’s new in psychtoolbox-3. Perception, 36(14), 1–16.

Kremer, G., & Baroni, M. (2011). A set of semantic norms for German and Italian. Behavior Research Methods, 43(1), 97–109. https://doi.org/10.3758/s13428-010-0028-x

Lesourd, M., Reynaud, E., Navarro, J., Gaujoux, V., Faye-Védrines, A., Alexandre, B., … Osiurak, F. (2023). Involvement of the posterior tool processing network during explicit retrieval of action tool and semantic tool knowledge: an fMRI study. Cerebral Cortex, 1–17. https://doi.org/10.1093/cercor/bhac522

Mahon, B. Z., & Caramazza, A. (2003). Constraining questions about the organisation and representationof conceptual knowledge. Cognitive Neuropsychology, 20(3–6), 433–450. https://doi.org/10.1080/02643290342000014

Marques, J. F., Raposo, A., & Almeida, J. (2013). Structural processing and category-specific deficits. Cortex, 49(1), 266–275. https://doi.org/10.1016/j.cortex.2011.10.006

Mathôt, S., Schreij, D., & Theeuwes, J. (2012). OpenSesame: An open-source, graphical experiment builder for the social sciences. Behavior Research Methods, 44(2), 314–324. https://doi.org/10.3758/s13428-011-0168-7

McRae, K., Cree, G. S., Seidenberg, M. S., & McNorgan, C. (2005). Semantic feature production norms for a large set of living and nonliving things. Behavior Research Methods, 37(4), 547–559. https://doi.org/10.3758/BF03192726

McRae, K., De Sa, V. R., & Seidenberg, M. S. (1997). On the Nature and Scope of Featural Representations of Word Meaning. Journal of Experimental Psychology: General, 126(2), 99–130. https://doi.org/10.1037/0096-3445.126.2.99

Montefinese, M., Ambrosini, E., Fairfield, B., & Mammarella, N. (2013). Semantic memory: A feature-based analysis and new norms for Italian. Behavior Research Methods, 45(2), 440–461. https://doi.org/10.3758/s13428-012-0263-4

Moss, H. E., & Tyler, L. K. (2000). A progressive category-specific semantic deficit for non-living things. Neuropsychologia, 38, 60–82.

Myung, J., Blumstein, S. E., & Sedivy, J. C. (2006). Playing on the typewriter, typing on the piano: Manipulation knowledge of objects. Cognition, 98(3), 223–243. https://doi.org/10.1016/j.cognition.2004.11.010

Myung, J., Blumstein, S. E., Yee, E., Sedivy, J. C., Thompson-Schill, S. L., & Buxbaum, L. J. (2010). Impaired access to manipulation features in Apraxia: Evidence from eyetracking and semantic judgment tasks. Brain and Language, 112(2), 101–112. https://doi.org/10.1016/j.bandl.2009.12.003

Peelen, M. V., & Caramazza, A. (2012). Conceptual object representations in human anterior temporal cortex. Journal of Neuroscience, 32(45), 15728–15736. https://doi.org/10.1523/JNEUROSCI.1953-12.2012

Perri, R., Zannino, G., Caltagirone, C., & Carlesimo, G. A. (2012). Alzheimer’s disease and semantic deficits: A feature-listing study. Neuropsychology, 26(5), 652–663. https://doi.org/10.1037/a0029302

Taylor, K. I., Devereux, B. J., & Tyler, L. K. (2011). Conceptual structure: Towards an integrated neurocognitive account. Language and Cognitive Processes, 26(9), 1368–1401. https://doi.org/10.1080/01690965.2011.568227

Tyler, L. K., Moss, H. E., Durrant-Peatfield, M. R., & Levy, J. P. (2000). Conceptual structure and the structure of concepts: A distributed account of category-specific deficits. Brain and Language, 75(2), 195–231. https://doi.org/10.1006/brln.2000.2353

Valério, D., Santana, I., Aguiar de Sousa, D., Schu, G., Leal, G., Pavão Martins, I., & Almeida, J. (2021). Knowing how to do it or doing it? A double dissociation between tool-gesture production and tool-gesture knowledge. Cortex, 141, 449–464. https://doi.org/10.1016/j.cortex.2021.05.008

Vivas, J., Vivas, L., Comesaña, A., Coni, A. G., & Vorano, A. (2017). Spanish semantic feature production norms for 400 concrete concepts. Behavior Research Methods, 49(3), 1095–1106. https://doi.org/10.3758/s13428-016-0777-2

Vivas, L., Montefinese, M., Bolognesi, M., & Vivas, J. (2020). Core features: measures and characterization for different languages. Cognitive Processing, 21(4), 651–667. https://doi.org/10.1007/s10339-020-00969-5

Warrington, E. W., & Shallice, T. (1984). Category specific semantic impairment. Brain, 107, 829–853.

